# Criticality between cortical states

**DOI:** 10.1101/454934

**Authors:** Antonio J. Fontenele, Nivaldo A. P. de Vasconcelos, Thaís Feliciano, Leandro A. A. Aguiar, Carina Soares-Cunha, Bárbara Coimbra, Leonardo Dalla Porta, Sidarta Ribeiro, Ana João Rodrigues, Nuno Sousa, Pedro V. Carelli, Mauro Copelli

**Affiliations:** Physics Department, Federal University of Pernambuco (UFPE), Recife, Brazil; Department of Biomedical Engineering, Federal University of Pernambuco, Recife, Brazil; Life and Health Sciences Research Institute (ICVS), School of Medicine, University of Minho, Braga, 4710-057, Portugal; ICVS/3Bs - PT Government Associate Laboratory, Braga/Guimarães, Portugal; Departamento de Morfologia e Fisiologia Animal, Universidade Federal Rural de Pernambuco (UFRPE), Recife, Brazil; Systems Neuroscience, Institut dInvestigacions Biomèdiques August Pi i Sunyer (IDIBAPS), Barcelona, Spain; Brain Institute, Federal University of Rio Grande do Norte (UFRN), Natal, RN, 59056-450, Brazil

**Author notes:** AJF and NAPV contributed equally.

## Abstract

Since the first measurements of neuronal avalanches [1], the critical brain hypothesis has gained traction [2]. However, if the brain is critical, what is the phase transition? For several decades it has been known that the cerebral cortex operates in a diversity of regimes [3], ranging from highly synchronous states (e.g. slow wave sleep [4], with higher spiking variability) to desynchronized states (e.g. alert waking [5], with lower spiking variability). Here, using independent signatures of criticality, we show that a phase transition occurs in an intermediate value of spiking variability. The critical exponents point to a universality class different from mean-field directed percolation (MF-DP). Importantly, as the cortex hovers around this critical point [6], it follows a linear relation between the avalanche exponents that encompasses previous experimental results from different setups [7, 8] and is reproduced by a model.

---

It is well established that cortical activity exhibits a rich repertoire of dynamical states [9–11]. This knowledge, initially based on electroencephalographic (EEG) recordings, later reached the spiking activity of large neuronal populations, in which the variability level has been used as a proxy of the cortical state [3, 12–14]. However, only recently has the diversity of cortical states been systematically considered in studies of criticality [15, 16].

In the first results that fuelled the critical brain hypothesis, Beggs and Plenz observed local field potential (LFP) neuronal avalanches *in vitro* with power law size distributions *P* (*s*) ~ *s*^−1.5^. The exponent coincides with that of a critical branching process, which has driven significant efforts towards the idea that the brain hovers around a critical point belonging to the mean-field directed percolation (MF-DP) universality class [17].

Findings for spiking data have, however, raised some controversy: on one hand, power law size distributions were found during strong slow LFP oscillations, under ketamine-xylazine [15] and isoflurane [16] anaesthesia, but on the other hand, long-range time correlations (another statistical signature of criticality [18]) were observed during fast LFP oscillations in freely-behaving rats, but not under ketamine-xylazine anaesthesia [15]. Those results lead to a conundrum, where the signatures of a critical state might be dependent on the level of synchronization, thus challenging the whole picture of directed percolation, which involves no oscillations whatsoever, and where the system goes from an absorbing to an active state.

To further investigate this topic we quantified the variety of cortical states in terms of the coefficient of variation (CV) of the summed population activity in the primary visual cortex of urethane-anaesthetized rats [14]. We recorded a total of 1628 units (Methods, Tables S1 and S2) in deep layers of the primary sensory cortex of urethane-anesthetized rats (*n* = 8), under spontaneous activity, during long periods (≥ 200 min). We employed high-count sites silicon probes (64/32 channels, see Methods), to record spiking activity of large neuronal populations (Fig. 1**a**). Given that the *in vivo* cortical dynamics crosses a wide range of states [19], the aim of this paper is to study the signatures of criticality as a function of the cortical state assessed by CV.

**FIG. 1.**
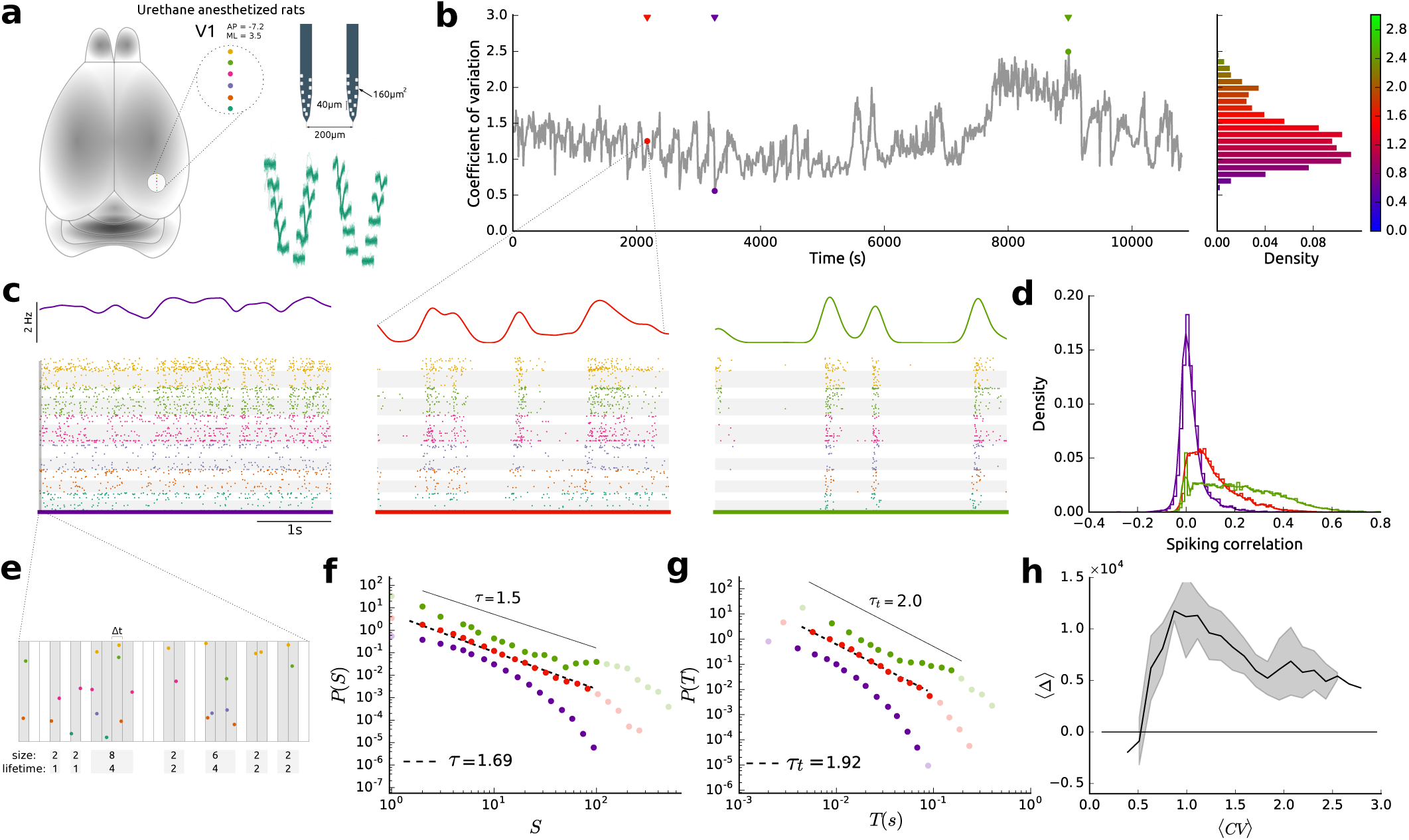
Statistical properties of cortical dynamics along different levels of spiking variability. **a**, Whole rat brain highlighting the position of the six shanks (colored dots) in the primary visual cortex (V1). Bottom right: samples of spike waveforms. **b**, (left) Coefficient of variation (CV) of the spiking activity in V1 (see also Figs. S2 and S3); each point was calculated for a 10s-long non-overlapping time period; colored marks (purple/red/green) indicate the level of spiking variability of three representative examples (low/intermediate/high, respectively); (right) CV histogram of a single animal. The vertical color bar is used as a reference for CV scale along the paper. **c**, Samples of 4s-long spiking activity across the three levels of spiking variability depicted in **b**; (up) population rate smoothed by a Gaussian kernel *σ* = 0.1 s; (bottom) raster plot: spikes recorded in the same shank are plotted with the same color shown in **a**; single/multi units (#SUA=138 and #MUA=153) with white/gray backgrounds, respectively. **d**, Histogram of pairwise spiking correlation along low/intermediate/high levels (same color as in **c**) where mean and standard deviation were 0.028 ± 0.069, 0.117 ± 0.115 and 0.224 ± 0.165, respectively (see Methods). **e**, The data is divided in non-overlapping time bins, Δt (see Methods). Population spikes preceded and followed by silences define a spike avalanche (gray backgrounds). The number of spikes defines the avalanche size, whereas the number of bins define its lifetime. **f,g**, Distributions of size and lifetime of spiking avalanches, *P* (*S*) and *P* (*T*) respectively, during the dynamical states described in **d**. Dark colored dots indicate the range to which we fitted a power-law in both cases. Black lines show the exponents of the MF-DP universality class. **h**, Relative goodness-of-fit test of the size distribution according to the Akaike Information Criterion (group data B, see Methods). Positive (negative) values indicate power-law (log-normal) behaviour.

Spiking cortical activity continuously changes its variability level under urethane anesthesia as well as in awake animals [12, 14]. The coefficient of variation (CV, Methods) has been used as an index of population spiking variability [12, 14]. We calculated CV within non-overlapping 10s windows. In that time scale, CV typically changes rapidly [3, 12, 14, 19] (Fig. 1**b**). Within each of these windows, for increasing values of CV, spiking activity ranges from completely desynchronized to a strongly synchronized state (Fig 1**c**). In what follows, we sort results according to CV values and average over consecutive percentiles to obtain 〈*CV*〉 as a representative of a given spiking variability level (Fig. S1). Different spiking correlation structures underlie those different regimes (Fig. 1**d**, *p* ≪ 0.01, ranksum test), in which both mean and standard deviation increase from the desynchronized state to the strongly synchronized state.

By dividing each 10s window in short time bins (Δ*t* ~ 2 − 4 ms), spike avalanches are defined by the spatio-temporal spiking patterns in between silent bins (Fig. 1**e**). We used the standard definition of Δ*t* as the average inter-spike interval [15, 20]. Since firing rates decrease monotonously with increasing 〈*CV* 〉 [21] (Fig. S4), for each 10s window a different Δ*t* was calculated. The size *S* and lifetime *T* of an avalanche are respectively given by the total numbers of spikes and bins within each event.

Even when bins are adjusted by firing rates, the statistics of avalanche size and duration differ across the range of 〈*CV* 〉 values. For instance, sampling the lower, intermediate and upper third portions of the 〈*CV* 〉 range, the degree to which the distributions of avalanche size and duration can be fitted by power laws (Methods)
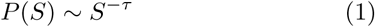

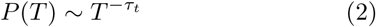

vary considerably (Fig. 1**f****,g**). In particular, the exponents *τ* and *τ_t_* do not necessarily agree with those originally obtained by Beggs and Plenz [20] for LFP avalanches in cultured slices (continuous lines Fig. 1**f****,g**). According to the Akaike criterion (Fig. 1**h**, Methods), power laws cease to be the best fitting distribution for sufficiently low 〈*CV* 〉 (and, if data is shuffled, for any 〈*CV* 〉, Figs. S5 and S6). Subsequently we refine our analysis by checking the consistency of scaling relations across 〈*CV* 〉 values.

The theory of critical phenomena predicts that, if Eqs. 1 and 2 hold at a critical point, then so does
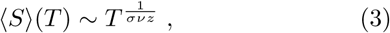

where 1/(*σνz*) is a combination of critical exponents [7, 22]. Our data is clearly consistent with Eq. 3, with the exponent 1/(*σνz*) depending on the spiking variability 〈*CV* 〉 (Fig. 2**a**). This scaling relation, however, is known to hold even far from criticality [7], so it can hardly be considered a sufficient signature in itself.

**FIG. 2.**
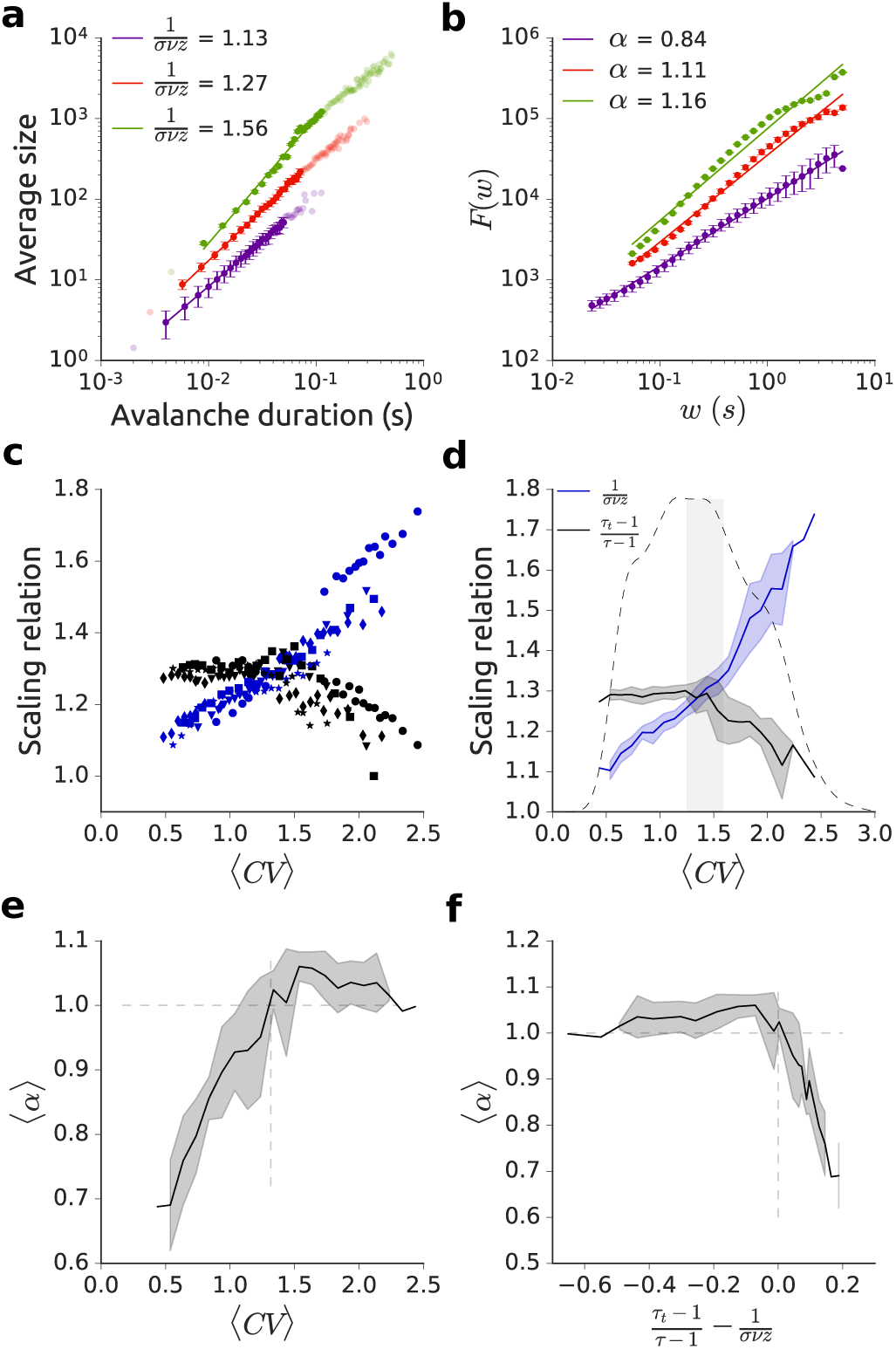
Signatures of criticality as a function of the cortical spiking variability. **a**, Power-law relation between size and lifetime of spikes avalanches across different levels of spiking variability (same color code as Fig. 1b). **b**, Root-mean squared fluctuation *F* of the detrended time series of the firing rates, versus window width *w*, across different levels of spiking variability (color code as Fig. 1b). **c**, Critical exponents relation across the variability spectrum per animal. Each symbol represents an animal. **d**, Group data of critical exponents relation across the variability spectrum. In both **c** and **d**, black (blue) corresponds to the left-hand (right-hand) side of Eq. 4. The dashed black curve represents the relative residence time across CV values. Gray stripe represents the critical value of 〈*CV* 〉∗ (1.4 ±0.2) where Eq. 4 holds. **e**, Group data of the DFA exponent *α* estimated along the variability spectrum. **f**, Group data of the DFA exponent *α* as a function of the difference between the scaling properties in **d**. (**c**–**f** : group data B, similar results for group data R in Fig S7).

If the system is indeed critical, the scaling relation between the lifetimes and sizes of avalanches (Eq. 3) is connected with the exponents governing their distributions (Eqs. 1 and 2), namely [7]:
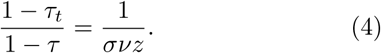

Since both sides of the above relation can be independently evaluated, we tested whether they equal each other as the brain spontaneously traverses the different levels of spiking variability. We verified that Eq. 4 clearly holds for *each* animal (*n* = 8) and, strikingly, the crossing between the left and right sides of the equation occurs around the same 〈*CV* 〉 value (Fig. 2**c**).

Averaging over animals (Fig. 2**d**), we obtain a critical value of spiking variability 〈*CV* 〉_∗_ = 1.4 ± 0.2, therefore far from the extremes of the variability spectrum. Moreover, when we compute the residence time distribution along this spectrum, we observe that the system spends most of the time close to the critical region (Fig. 2**d**). This is consistent with a scenario in which the urethanized brain hovers around a critical point [6]. We found similar results in a different strain (non-albino rats, Long Evans, *n* = 3), using a 20% lower spatial resolution (8 sites/shank) (Methods and Figs. S6 and S7).

Another feature of systems at the critical point is self-affinity of time series over different time scales, as assessed by detrended fluctuation analysis (DFA) [18, 25]. For our data, the root-mean squared fluctuation *F* scales with the length of the time window *w* as *F* ~ *w^α^* (Methods), and *α* also depends on 〈*CV* 〉 (Figs. 2**b**,**e**).

We found *α* ≈ 1, indicating long-range time correlations and 1/*f* noise, at the same critical value of 〈*CV* 〉 that was independently inferred by the scaling relation Eq. 4 (Fig. 2**e**). There is, therefore, a remarkable convergence between these two independent signatures of criticality (Fig. 2**f**).

If the dynamics of the urethane-anaesthetized brain indeed hovers around a critical point, then what are the avalanche exponents at criticality? At the critical value 〈*CV* 〉_∗_, the exponents are close to their minimal values and we obtain *τ* = 1.52 ± 0.09 and *τ_t_* = 1.7 ± 0.1 (Figs. 3**a** and 3**b**).

**FIG. 3.**
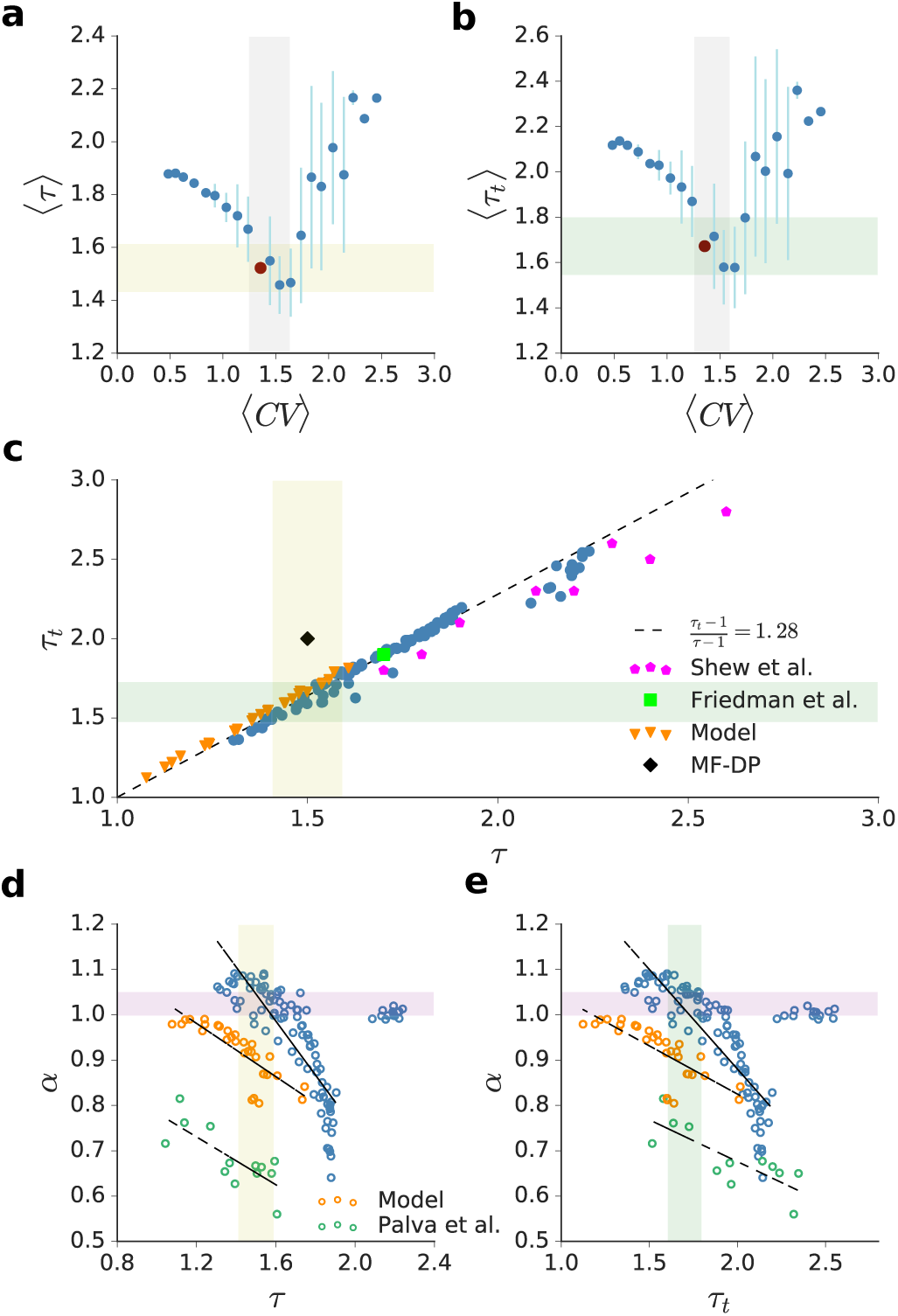
Group data of correlations between exponents governing long-range time correlations, size and life-time distributions. **a,b**, Exponents of the size and life-time distributions along the levels of variability. Gray stripe represents the critical value of 〈*CV* 〉∗ (within errors) where Eq. 4 holds. Yellow (green) stripe represents the value of 〈*τ* 〉 (〈*τt*〉) compatible with 〈*CV* 〉∗. **c**, Linear relation between the size and lifetime critical exponents across animals. We also plot other experimental results [7, 8] and those obtained by a model [23]. The black symbol corresponds to MF-DP values. **d,e**, Spread of DFA and avalanche exponents across animals displaying negative correlation (*r* = −0.85/*r* = −0.86). Similar results are obtained for M/EEG data in humans [24] (*r* = −0.77/*r* = −0.79) and a model [23] (*r* = −0.82/*r* = −0.78). Purple stripe represents the value of *α* compatible with 〈*CV* 〉∗. (**a**–**e**: group data B).

Note that while the value of *τ* coincides with the critical exponent of MF-DP [26], the value of *τ_t_* does not. The disagreement with the MF-DP universality class, regardless of the 〈*CV* 〉 value, is clearly seen when results are parametrically plotted in the (*τ, τ_t_*) plane (Fig. 3**c**). We find, moreover, that other results in the literature (from diverse experimental recordings such as the *ex-vivo* visual cortex of the turtle [8] and *in vitro* cultured slices of the rat cortex [7]) lie close to the linear spread of the avalanche exponents of our data (Fig. 3**c**).

Such a coincidence suggests a common underlying mechanism. A similar linear trend is found in the CROS (“CRitical OScillations” [27]) model with excitatory and inhibitory neurons in which a transition occurs at the onset of collective oscillations [23, 27] (Fig. 3**c**). The model successfully mimics our experimental results, in the sense that it also presents DFA exponents close to one at criticality and avalanche exponents vary continuously near the transition [23]. This supports a scenario in which the transition governing brain dynamics is not between absorbing and active phases, but rather between active and oscillating phases [27–29].

Finally, it is interesting to delve deeper into the relation between the DFA exponent *α* and the pair of avalanche exponents *τ* and *τ_t_* (Fig. 2**f**). In principle they pertain to very different time scales [24]. Yet, as 〈*CV* 〉 varies, these two very different fingerprints of criticality keep a negative correlation (Figs. 3**d** and 3**e**). This result is reminiscent of those obtained by Palva et al. for M/EEG data of human subjects [24]. Once more, the CROS model [23] shows a reasonable agreement with our experimental results (Fig. 3**d** and 3**e**), reinforcing the idea of a transition at the onset of oscillations.

In conclusion, we found consistent markers of criticality (scaling in avalanche statistics as well as long-range time correlations) in the cortical activity of urethane-anaesthetized rats. The critical point is neither at the synchronous nor the asynchronous ends of the spectrum, but rather at an intermediate value 〈*CV* 〉_∗_ of the coefficient of variation. Those results hold for group data, but are also verified at each subject (*n* = 8).

Importantly, our results are incompatible with a DP-like phase transition between a quiescent and an active state, a paradigm which has been a *de facto* theoretical workhorse of the field for over a decade [17]. We found a linear relationship between *τ* and *τ_t_* across cortical states that encompasses results from other experimental setups and is reproduced by the CROS model.

These results open new experimental as well as theoretical avenues. On one hand, the present results can guide further development of models for criticality in the brain. On the other hand, it remains to be investigated whether densely recorded activity in awake animals will yield similar results. Moreover, the present results highlight the relevance of intermediate levels of spiking variability for state-dependent processing in the primary sensory cortex. We propose that, if the cortex demands both modes of operation (synchronized and desynchronized) for different functions [30], it might be advantageous to self-organize near and hover over the critical point between them.

## a. Acknowledgments

C.S.-C. was recipient of the Fundação para a Ciência e Tecnologia (FCT) Fellowship SFRH/BD/51992/2012 and is currently recipient of a post-doctoral fellowship from the Programa de Atividades Conjuntas (PAC), through MEDPERSYST Project POCI-01-0145-FEDER-016428 (supported by the Portugal2020 Programme). B.C. is recipient of a PhD scholarship funded by FCT (SFRH/BD/98675/2013). A.J.R. is a FCT Investigator Fellow (IF/00883/2013). N.A.P.V. was a recipient of CNPq Grant 249991/2013-6 and CAPES Grant 88887.131435/2016-00. This work was developed under the scope of the project NORTE-01-0145-FEDER-000013, supported by the Northern Portugal Regional Operational Programme (NORTE 2020), under the Portugal 2020 Partnership Agreement, through the European Regional Development Fund (FEDER). Part of the work was supported by the Janssen Neuroscience Prize (1st edition) and by the BIAL Grant 30/2016. A.J.F., T.F. and L.A.A.A. acknowledge support from CAPES, FACEPE and CNPq. S.R. benefits from CNPq (grants 308775/2015-5 and 408145/2016-1). L.D.P. and M.C. were supported by FACEPE and CAPES. P.V.C. was supported by FACEPE (grant APQ 0826-1.05/15) and CAPES. M.C. is also supported by CNPq (grant 310712/2014-9). S.R. and M.C. are also supported by Fundação de Amparo à Pesquisa do Estado de São Paulo Grant #2013/07699-0 (Center for Neuromathematics).

## I. METHODS

### A. Group data B

Part of this dataset was previously described [14]. We used Wistar-Han rats (*n* = 5, male, 350-500 g, 3-6 months old, Charles River) in our recordings. We anaesthetized the animals with 1.44 g/kg of fresh urethane, diluted at 20% in saline, in 3 injections (i.p.), 15 min apart. In some animals a supplement has been administrated in order to reach the proper levels of anaesthesia. We placed the rats in a stereotaxic frame and marked, based on the Paxinos Atlas [31], the coordinates to access V1 (Bregma: AP = -7.2, ML = 3.5). A cranial window (2.5 mm diameter) was opened at this site. We recorded extra-cellular voltages using a 64-channels silicon probe (BuzsakiA64sp, Neuronexus) with 60 sites along 6 shanks, 10 sites/shank with impedance of 1–3 MOhm at 1 kHz; shanks were 200 *µ*m apart and the area of each site was 160 *µ*m^2^, disposed from the tip in a staggered configuration, 20 *µ*m apart (Fig. 1**a**). All data from the silicon probe were sampled (30 kHz), amplified and digitized in a single head-stage (Intan RHD2164). We used the Klusta-Team [32, 33] software to perform spike sorting on raw electrophysiological data, running in two computational infrastructures (LFTC/UFPE and NPAD/UFRN). Health monitoring of animals was performed according to FELASA guidelines. All manipulations were conducted in strict accordance with European Regulations (European Union Directive 2010/63/EU). Animal facilities and the people directly involved in animal experiments were certified by the Portuguese regulatory entity DGAV. All the experiments were approved by the Ethics Committee of the University of Minho (SECVS protocol #107/2015). The experiments were also authorized by the national competent entity DGAV (#19074).

### B. Group data R

We recorded a second dataset to test the robustness of the results in three different ways: 1) experiments performed in a different laboratory 2) with a different non-albino strain and 3) fewer and less dense recording sites. This data set used Long-Evans rats (*n* = 3, male, 250-360 g, 3-4 months old) in our recordings. We anaesthetised the animals with 1.58 g/kg of fresh urethane, diluted at 20% in saline, in 3 injections (i.p.), 15 min apart. Stereotaxic positioning was performed as in group B. We recorded extra-cellular voltages using a 32-channels silicon probe (BuzsakiA32, Neuronexus) with 32 sites along 4 shanks, 8 sites/shank with impedance of 1–3 MOhm at 1 kHz; shanks were 200 *µ*m apart and each sites area was 160 *µ*m^2^, disposed from the tip in a staggered configuration, 20 *µ*m apart. All data from the silicon probe were sampled (24 kHz), amplified and digitized in a PZ2 TDT, that transmits to a RZ2 TDT base station. Spike sorting was performed as in group B. Housing, surgical and recording procedures were in strict accordance with the CONCEA - MCTI, and was approved by the Federal University of Pernambuco (UFPE) Committee for Ethics in Animal Experimentation (23076.030111/2013-95 and 12/2015).

### C. Coefficient of variation

We evaluate the global firing rate *r*(*t*) in a time window Δ*T* using the spikes of every recorded unit
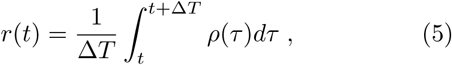

where the neural response is 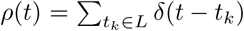 and Δ*T* = 50 ms. The coefficient of variation *CV* was calculated for each non-overlapping 10-second window (Figs. 1**b** and S1)
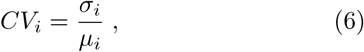

where *σ* and *µ* are the standard deviation and mean of the firing rate in the *i*-th 10-second window (Fig. S1). Each dataset had 1080 *CV* values.

### D. Spiking correlations

The firing rate *r_i_*(*t*) of the *i*-th spike train was calculated according to Eq. 5 with Δ*t* = 1 ms, *r_i_*(*t*) ∊ {0, 1} ms^−1^. The time series *n_i_*(*t*) was obtained by the convolution of *r_i_*(*t*) with a kernel *h*,
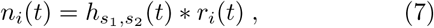

where *h* is a Mexican-hat kernel which was obtained by the difference between the zero-mean Gaussians with standard deviations *s*_1_ = 100 ms and *s*_2_ = 400 ms [12].

The spiking correlation coefficient between two units *i* and *j* is given by:
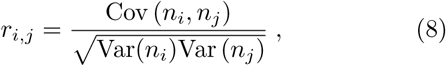

where Var and Cov are the variance and covariance, respectively.

### E. Critical exponents estimation

The exponents governing power-law distributions were obtained via a Maximum Likelihood Estimator procedure [34, 35]. To fit a discrete power-law distribution
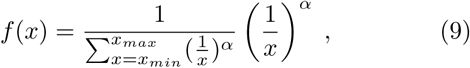

the lower bound *x_min_* was set to the smallest observable (size, lifetime). For dataset B (R), the upper bound *x_max_* was set to 100 (40) for size distributions and 25 (15) time bins for lifetime distributions. Next, we estimate *α* by maximizing the likelihood function
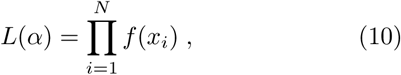

where *N* is the number of measurements. We work with the much more convenient log-likelihood
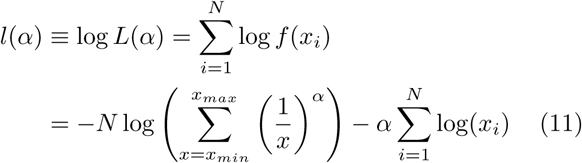

and obtain 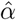, the parameter that best fits a power-law, by
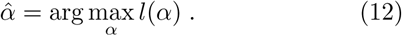

The maximization of *l*(*α*) is obtained via a lattice search algorithm [36].

### F. Relative goodness of fit test

We used the Akaike information criterion (AIC) as a measure of the relative quality of a given statistical model for a data set [37]:
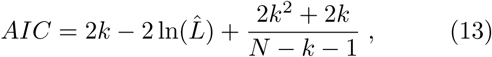

where 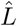, is the likelihood at its maximum, *k* is number of parameters and *N* the sample size. The lower the AIC, the better the model fits the data. We defined Δ = *AIC_ln_*−*AIC_pl_*, where *AIC_ln_* and *AIC_pl_* correspond to the AIC of a log-normal and a power-law model, respectively. Therefore, Δ ≥ 0 (Δ ≤ 0) implies that the power-law model is better (worse) than the log-normal to fit the data.

### G. Detrended Fluctuation Analysis

We used detrended fluctuation analysis (DFA) to assess scale-invariant long-range time correlations [15, 25]. Given a time series, *z*, and a width of time window, *w*: (1) obtain an integrated zero-mean version from the original time series 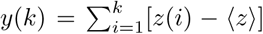; (2) divide the whole time period in consecutive non-overlapping bins of width *w* and fit the local linear trend, *y_w_*, with the least squares methods within each bin; (3) then, the fluctuation at a given time-scale *w* is calculated with respect to the linear trend by
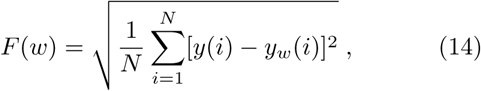

where *N* is number of consecutive non-overlapping time windows.

### H. Computational model

We made use of the CROS model previously described in Refs. [23, 27]. Briefly, the model consists of a two-dimensional 300 × 300 network of excitatory (75%) and inhibitory (25%) stochastic integrate-and-fire neurons. Each neuron is locally connected to its neighbors within a square of size 7 × 7 centered around it. Connectivity is controlled by two free parameters of this model, namely the fractions of excitatory and inhibitory outgoing synapses of the total number of neurons within the local range. These parameters vary within the ranges 2 − 60% and 30 − 90%, respectively. The model has no absorbing state because excitatory neurons receive a low constant Poisson input. In this model each avalanche is initiated when the activity of the network (defined as the total sum of spike activity) crosses a threshold (defined here as 30% of the median spike activity of the network). The duration of an avalanche is given by the total time in which the spike activity of the network remains above the threshold. Detrended fluctuation analysis is calculated directly on the global network activity. The model exhibits signatures of criticality at the onset of collective oscillations, where a peak emerges in the Fourier spectrum of the network activity and DFA exponent approaches unity. For details of the model parameters and implementation, see reference [23].

